# A method for the generation of pseudotyped virus particles bearing SARS coronavirus spike protein in high yields

**DOI:** 10.1101/2021.07.30.454063

**Authors:** Yoichiro Fujioka, Sayaka Kashiwagi, Aiko Yoshida, Aya O. Satoh, Mari Fujioka, Maho Amano, Yohei Yamauchi, Yusuke Ohba

## Abstract

The ongoing severe acute respiratory syndrome coronavirus 2 (SARS-CoV-2) pandemic has threatened human health and the global economy. Development of additional vaccines and therapeutics is urgently required, but such development with live virus must be conducted with biosafety level 3 confinement. Pseudotyped viruses have been widely adopted for studies of virus entry and pharmaceutical development to overcome this restriction. Here we describe a modified protocol to generate vesicular stomatitis virus (VSV) pseudotyped with SARS-CoV or SARS-CoV-2 Spike protein in high yield. We found that pseudovirions produced with the conventional transient expression system lacked coronavirus Spike protein at their surface as a result of inhibition of parental VSV infection by overexpression of this protein. Establishment of stable cell lines with an optimal expression level of coronavirus Spike protein allowed the efficient production of progeny pseudoviruses decorated with Spike protein. This improved VSV pseudovirus production method should facilitate studies of coronavirus entry and development of antiviral agents.

## INTRODUCTION

Severe acute respiratory syndrome coronavirus 2 (SARS-CoV-2) is an enveloped, positive-strand RNA virus that belongs to the *Coronaviridae* family and is responsible for the recent pandemic of coronavirus disease 2019 (COVID-19). As of 15 June 2021, there had been >174 million confirmed cases of and ∼3.7 million deaths from COVID-19 reported in over 220 countries, areas, or territories (https://www.who.int/emergencies/diseases/novel-coronavirus-2019).

The envelope of SARS-CoV-2, like that of SARS-CoV, contains three structural proteins: the spike glycoprotein (hereafter referred to simply as S protein), membrane protein, and small envelope protein (Schoeman and Fielding, 2019; Wrapp et al., 2020). Together with retrovirus envelope proteins and influenza virus hemagglutinin proteins, S protein is categorized as a class I viral fusion protein (Millet and Whittaker, 2018; White 2008), and it is targeted to the rough endoplasmic reticulum (rER) of host cells by an NH_2_-terminal signal sequence (Breitling and Aebi, 2013; Duan et al., 2020; Lontok et al., 2004; Sadasivan et al., 2017). Cleavage of the signal sequence is followed by resumption of peptide chain elongation and insertion of the synthesized S protein into the ER membrane as a homotrimer (Binns et al., 1985; Braakman and Hebert, 2013; Delmas and Laude, 1990; Duan et al., 2020). S protein plays a key role in both specific interaction of the virus with its host cell receptor and subsequent virus internalization via endocytosis or membrane fusion. Binding of the coronavirus to its host cell receptor, angiotensin-converting enzyme 2 (ACE2), has been attributed to the S1 receptor-binding subunit region of S protein, which adopts an active “standing-up” conformation for receptor binding and an inactive “lying-down” conformation for immune evasion (Guan et al., 2020; Wrapp et al., 2020). After the S1 subunit region binds to ACE2, S protein undergoes proteolytic cleavage into S1 and S2 fusion subunits at the S1/S2 site in the presence of transmembrane protease serine 2 (TMPRSS2) and furin (Bestle et al., 2020; Jaimes et al., 2020). Additional cleavage sites that are located within the S2 subunit and are cleaved by host cell proteases including cathepsins, furin-like proprotein convertases, and trypsin-like serine protease (Burkard et al., 2014; Coutard et al., 2020; Hoffmann et al., 2020). The exposed S2 subunit mediates membrane fusion either at the plasma membrane or with the endosomal membrane, depending on protease availability in the host cell. Characterization of S protein is thus crucial for a better understanding of SARS-CoV-2 entry, but such studies are often limited because experiments with the live virus must be conducted in a biosafety level 3 (BSL-3) containment facility.

Pseudotyped virus systems are highly useful for characterization of viral envelope proteins and investigation of their role in virus entry (Hoffmann et al., 2020; Miyauchi et al., 2009; Nanbo et al., 2010; Rentsch and Zimmer, 2011; Schelhaas et al., 2012; de Vries et al., 2011). Virus particles produced with such a system consist of a surrogate viral core and heterologous viral envelope proteins. The genome of the parental virus is modified to remove essential genes for viral reproduction, so as to prevent the generation of infectious progeny viruses. This approach allows viruses pseudotyped with envelope proteins of highly pathogenic viruses to be handled safely in a BSL-2 containment facility instead of a BSL-3 or BSL-4 facility. Vesicular stomatitis virus (VSV), a negative-strand RNA virus of the *Rhabdoviridae* family, has been widely adopted in both pseudotyped and recombinant virus systems (Hoffmann et al., 2020; Miyauchi et al., 2009; Nanbo et al., 2010; Rentsch and Zimmer, 2011). It possesses five structural proteins—the glycoprotein (G), large polymerase protein (L), matrix protein (M), nucleocapsid (N), and phosphoprotein (P) (Lichty et al., 2004)—among which G protein contributes to binding to the host cell surface and fusion with the endosomal membrane. VSV was the first negative-strand RNA virus for which a reverse genetics approach was established (Lawson et al., 1995), an approach that allowed packaging of a reporter gene into the viral genome for quantitative evaluation of infectivity of recombinant viruses on the basis of the reporter gene activity. It was also found that a large amount of morphologically intact virus particles could be produced even in the absence of G protein. In addition, coinfection of cells with VSV and other viruses results in the formation of VSV pseudotyped with heterologous envelope proteins (Huang et al., 1974; Witte and Baltimore, 1977). With these features, VSV-based pseudotyping is a potentially powerful tool for studies of the role of envelope proteins, including S proteins of SARS-CoV and SARS-CoV-2, in virus entry (Clausen et al., 2020; Hoffmann et al., 2020).

During the course of experiments with VSV pseudoviruses bearing S protein of SARS-CoV or SARS CoV-2 described here, we found that most pseudotyped particles produced by the conventional transfection protocol lack S protein, likely because S protein expression at a high level in the virus-producing cells suppresses infection by parental VSV. We therefore developed an improved protocol based on cell lines stably expressing S proteins. This new method generates VSV pseudotyped particles decorated with S protein at high efficiency and with high infectious yields compared to the conventional approach. Our protocol increases the fidelity of SARS-CoV-2 entry research and should facilitate the development of new antiviral agents.

## RESULTS AND DISCUSSION

We initially aimed to evaluate the loading efficiency of S proteins on VSV pseudotyped virus particles. According to the published protocol (Rentsch and Zimmer, 2011), HEK293T cells transiently expressing SARS-CoV S protein were exposed to VSVΔG bearing G protein of VSV (VSVΔG-G), and the resulting culture supernatant was harvested as a pseudovirus suspension in the presence of neutralizing antibodies to VSV G protein (to eliminate parental VSVΔG-G). The pseudotyped virions obtained in this manner were stained with the lipophilic carbocyanine dye DiI and then subjected to immunofluorescence analysis with a rabbit monoclonal antibody to SARS-CoV S protein. Confocal fluorescence imaging revealed that only a small proportion of the obtained particles harbored S protein (**Figure 1A**).

**Figure 1.**
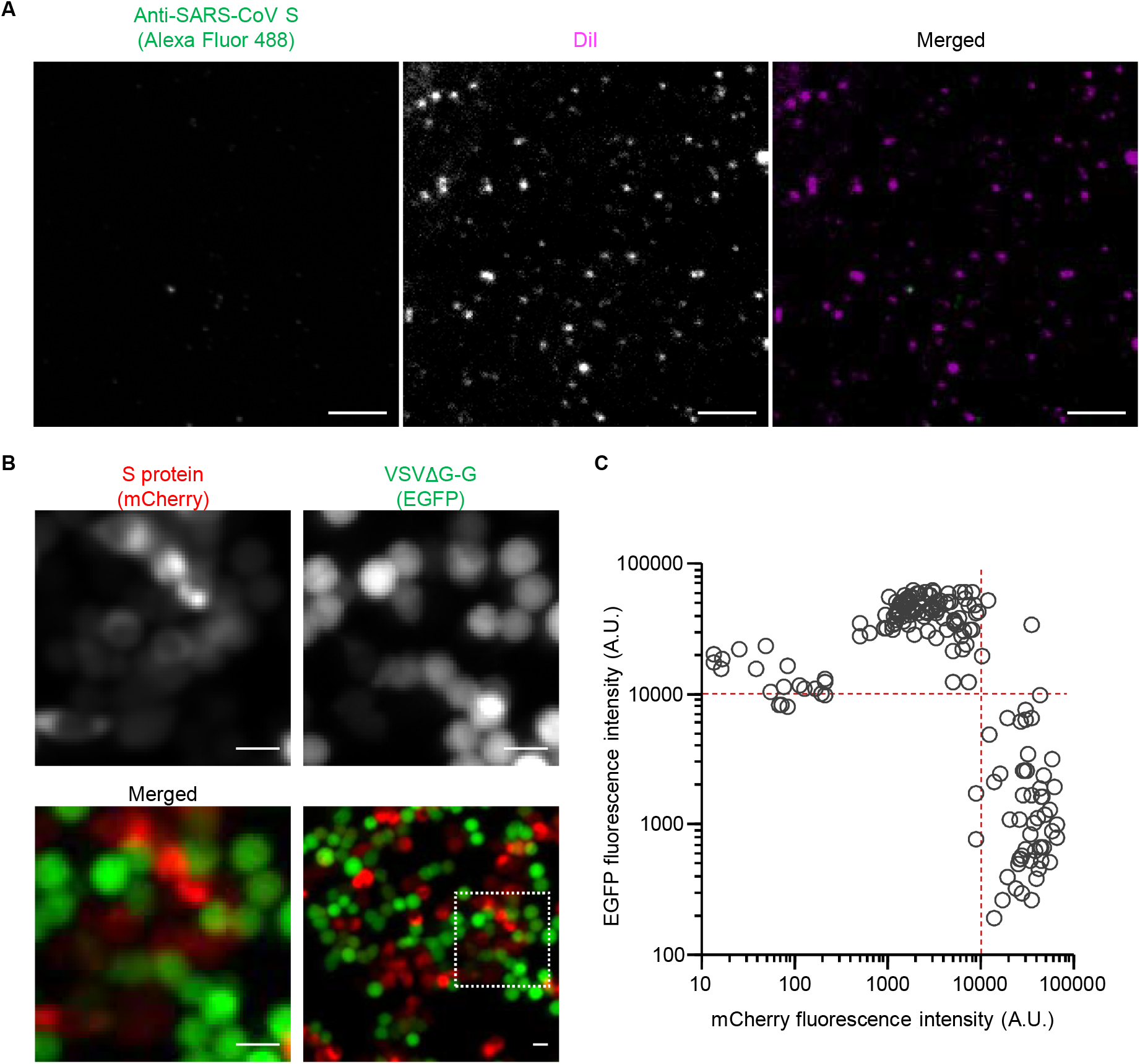
Pseudotyped virus particles produced from cells transiently expressing SARS-CoV S protein are not loaded with S protein. **(A)** HEK293T cells were transfected with an expression vector for SARS-CoV S protein as well as infected with VSVΔG-G. The culture supernatant was collected and stained with DiI as well as subjected to immunofluorescence staining with antibodies to SARS-CoV S protein and Alexa Fluor 488–conjugated secondary antibodies. The virus particles were then adsorbed onto a polyethylenimine-coated glass-based plate and observed with a confocal fluorescence microscope. Representative images are shown. Bars, 10 μm. **(B)** HEK293T cells were transfected with an expression vector for mCherry-tagged SARS-CoV S protein and infected with VSVΔG-G for 16 h. They were then observed with a fluorescence microscope for detection of mCherry (red) and GFP (green) fluorescence. Representative images are shown. The boxed region in the lower right panel is shown at higher magnification in the other panels. Bars, 10 μm. **(C)** Fluorescence intensities of GFP and mCherry in individual cells as in (**B**). See also **Figure S1**.

To examine S protein expression in the pseudotyped virion-producing cells, we prepared an expression vector for a fluorescent protein-tagged form of the protein. The coding sequence for mCherry, a red fluorescent protein (RFP), was thus inserted upstream of the NH_2_-terminal signal sequence of S protein, with the expectation that the fluorescent tag would be cleaved together with the signal sequence in the ER and so would not suppress virion formation. Indeed, immunoblot analysis of HEK293T cells transfected with the vector for mCherry-tagged S protein revealed separate fragments corresponding to mCherry and to S protein (**Figure S1A**), and no red fluorescence signal was detected from the VSVΔG-mCherry-S pseudoparticles produced by these cells (**Figure S1B**). Furthermore, determination of focus-forming units (FFU) by counting the number of cells positive for green fluorescent protein (GFP) encoded by the reporter gene packaged in the genome of parental VSVΔG (Rentsch and Zimmer, 2011) did not reveal a marked difference in infectivity between VSVΔG-mCherry-S and VSVΔG-S pseudoviruses in BEAS-2B cells expressing ACE2, also a receptor for SARS-CoV (**Figure S1C**). This result also supported the notion that the chimeric protein undergoes proteolytic cleavage and does not inhibit particle formation.

Fluorescence imaging of HEK293T cells transfected with expression vectors for mCherry-S or mCherry showed that mCherry-S expression interfered with VSVΔG-G infection; the GFP signal was thus detected in cells with no or only a low level of mCherry-S expression but not in highly red fluorescent cells (**Figure 1B**). Quantitative analysis revealed that GFP-positive cells were indeed highly enriched in cells in which red fluorescence intensity was <10,000 arbitrary units (A.U.) (**Figure 1C**), whereas a high level of mCherry expression only partially suppressed VSVΔG-G infection (**Figure S1D**). These observations also suggested that virus particles lacking envelope protein might be produced from cells with no mCherry-S expression. Indeed, immunofluorescence analysis with antibodies to VSV M protein showed that the supernatant of vector-transfected, VSVΔG-G-exposed HEK293T cells contained fluorescent puncta even when the vector did not encode an envelope protein (**Figure S1E**).

Our results together suggested that a low and persistent level of S protein expression might be required for the production of pseudotyped virus particles that uniformly bear S protein. We therefore attempted to establish cell lines that stably express S protein. However, although HEK293T cells are used in the standard protocol for VSV-based pseudotyped virus production, we found that HEK293T cells expressing mCherry-S manifested a shrunken morphology compared with those expressing mCherry alone (**Figure S1F**). We next introduced the expression vector for mCherry-S into VeroE6 cells, which have been widely adopted for SARS-CoV research. Fluorescence imaging showed that the morphology of VeroE6 cells expressing mCherry-S was similar to that of those expressing mCherry alone (**Figure 2A**). To generate stable cell lines, we isolated transfected VeroE6 cells by treatment with trypsin and subjected them to puromycin selection. After incubation for 24 h, multinucleated cells, indicative of cell-cell fusion resulting from S protein activation by trypsin cleavage, were apparent for the cells expressing mCherry-S (**Figure 2B**). To avoid cleavage of S protein, we used enzyme-free cell-dissociation solutions for cell isolation. Treatment with such solutions did not induce cleavage of S protein (**Figure S2A**) or multinucleated cells (**Figure 2C**), thus allowing the generation of cell lines stably expressing S protein (**Figure 2D**).

**Figure 2.**
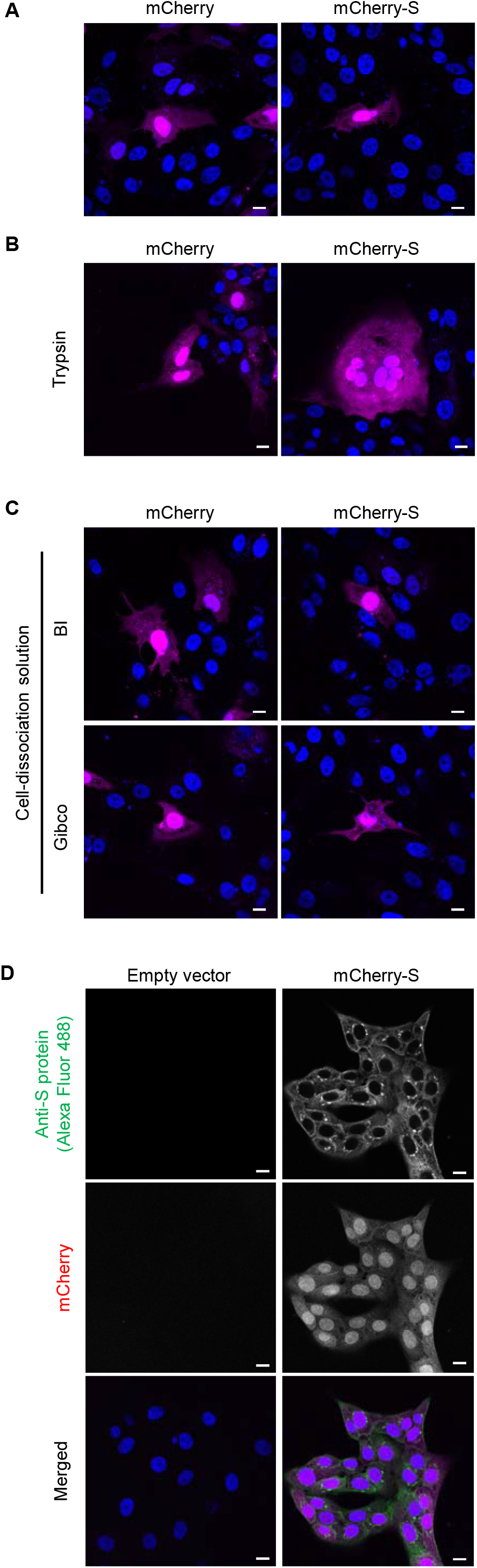
Trypsin treatment induces cell-cell fusion of S protein-expressing cells. (**A**) VeroE6 cells transfected with an expression vector for mCherry or mCherry-tagged S protein of SARS-CoV (mCherry-S) that had been linearized with *Puv*I were stained with Hoechst 33342 and observed by fluorescence microscopy. Representative images of mCherry (magenta) and Hoechst 33342 (blue) fluorescence are shown. Bars, 10 μm. (**B** and **C**) Cells prepared as in (**A**) were harvested by exposure to trypsin (**B**) or to enzyme-free cell dissociation solutions from Biological Industries (BI) or Gibco (**C**). They were then transferred to cell culture dishes and cultured for 24 h before fluorescence microscopic analysis as in (**A**). Representative images are shown. Bars, 10 μm. (**D**) Cells transfected with the mCherry-S vector (or the corresponding empty vector) as in (**A**) were cultured in the presence of puromycin for 14 days, stained with Hoechst 33342, and subjected to immunofluorescence analysis with antibodies to SARS-CoV S protein and Alexa Fluor 488–conjugated secondary antibodies. Representative fluorescence microscopic images of S protein immunostaining (green) as well as of mCherry (magenta) and Hoechst 33342 (blue) fluorescence are shown. Bars, 10 μm. See also **Figure S2**.

We next examined whether such a cell line was susceptible to VSVΔG-G infection for production of pseudoviruses harboring S protein. Fluorescence microscopy and quantitative analysis of the virus-exposed cells revealed that the mCherry fluorescence intensity of all cells was <10,000 A.U. (the desired upper limit as determined in **Figure 1C**), and that most cells were positive for GFP, indicative of uniform VSVΔG-G infection (**Figures S2B** and **S2C**). The virus particles produced by the cells were then stained with DiD and adsorbed on a polyethylenimine-coated glass-based dish to determine the amount of particles per unit volume by measurement of the area of DiD-positive puncta with a confocal microscope. The amount of particles produced by VeroE6 cells stably expressing mCherry-S was ∼10% of that of those produced by HEK293T cells transiently expressing mCherry-S (**Figures S3A** and **S3B**). BEAS-2B cells expressing ACE2 were then exposed to the pseudoviruses for 1 h at 4°C (to allow for virus attachment) and subjected to immunofluorescence analysis with antibodies to SARS-CoV S protein and to VSV M protein in order to allow visualization of S protein and all pseudovirus particles, respectively. Confocal imaging showed that, in contrast to the virus particles produced by transiently transfected HEK293T cells, essentially all virus particles produced by the stably transfected VeroE6 cell line were positive for S protein (**Figure 3A**). Indeed, quantitative analysis revealed that the fraction of S protein-positive particles generated by the stable cell line was about seven times that generated by the transiently transfected cells (**Figure 3B**). These results thus demonstrated the efficient production of pseudoviruses that were essentially all loaded with S protein by the stable cell line.

**Figure 3.**
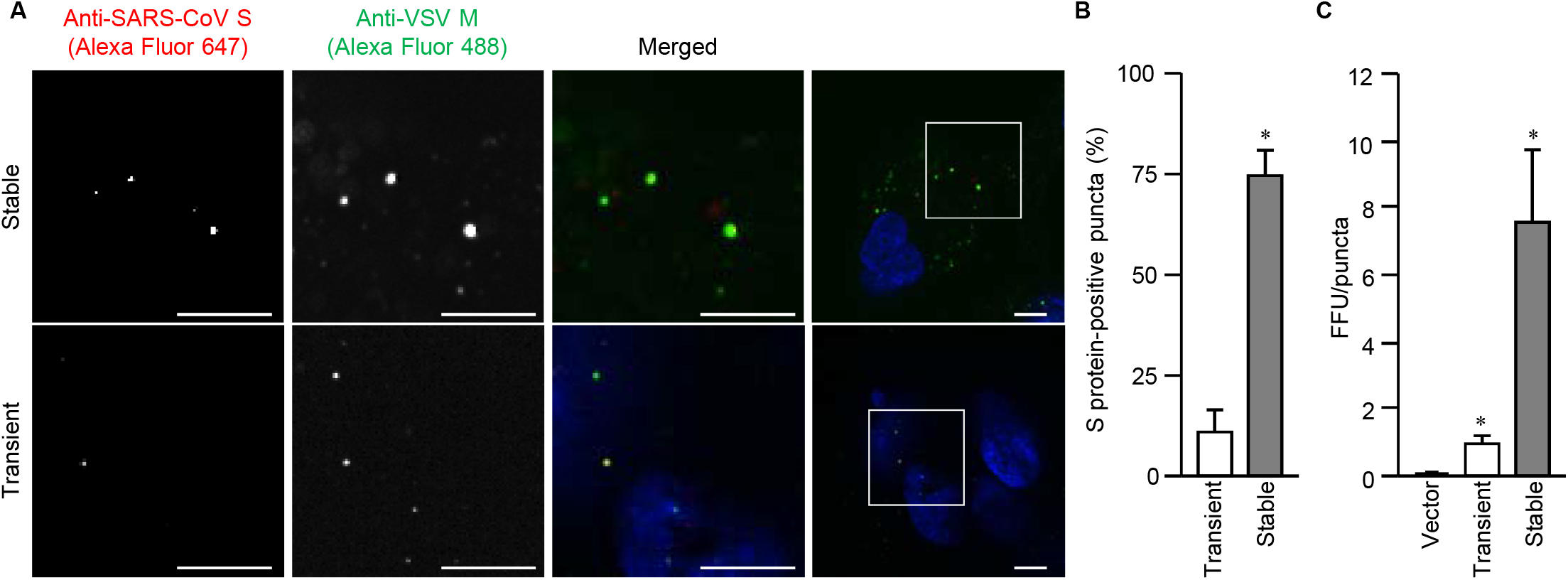
A large proportion of pseudotyped virus particles produced from VeroE6 cells stably expressing SARS-CoV S protein are loaded with S protein. **(A)** VeroE6 cells stably expressing or HEK293T cells transiently expressing SARS-CoV S protein were infected with VSVΔG-G for 16 h, and equal numbers of pseudoviruses released into the culture supernatants were added to ACE2-expressing BEAS-2B cells for 1 h at 4°C. The latter cells were then fixed, stained with Hoechst 33342 (blue), subjected to immunofluorescence analysis with antibodies to SARS-CoV S protein and VSV M protein, and observed with a confocal microscope. Representative images are shown. The boxed regions in the right panels are shown at higher magnification in the panels to the left. Bars, 10μm. **(B)** Quantification of the fraction of S protein-positive puncta among all M protein–positive puncta for images as in (**A**). Data are means + SEM from three independent experiments. *, *p* < 0.001 (Student’s *t* test). **(C)** BEAS-2B cells stably expressing ACE2 were exposed to pseudotyped viruses prepared as in (**A**) for determination of the number of GFP-positive cells and calculation of FFU of virus suspension. FFU was normalized by the amount of pseudotyped viruses, which was determined by the area of puncta of DiD-stained viruses on a polyethylenimine-coated glass-based plate. Data are means + SEM from three independent experiments. *, *p* < 0.004 (one-way ANOVA with Tukey’s HSD post hoc test). See also **Figure S3**.

We further examined whether the titer of pseudovirus particles generated by the stable cell line was higher than that of particles generated by the conventional method. There was no significant difference between the FFU values obtained for the two types of pseudoviruses (**Figure S3C**). To normalize infectivity, we prepared dilutions of the virus suspensions (down to 1.0 FFU/µl) and plotted FFU against the area of puncta. A linear relation was apparent for each type of pseudovirus, although the slopes of the regression lines differed (**Figure S3D**). The FFU value normalized by the amount of viruses produced by the stable cell line was 6.4 times the corresponding value for the viruses produced by the transiently transfected cells (**Figure 3C**), indicating that the modified method established in this study results in a substantial improvement in infectious pseudovirus particle production.

Finally, we attempted to produce pseudotyped virus particles with envelopes bearing the S protein of SARS-CoV-2 by our newly established protocol. We thus established a VeroE6 cell line that stably expresses SARS-CoV-2 S protein (**Figure S4A**). The fraction of SARS-CoV-2 S protein-positive puncta (**Figures 4A** and **4B**) and FFU normalized by the area of DiD-positive puncta (**Figures 4C, S4B** and **S4C**) were five and seven times as high, respectively, for viruses produced from the VeroE6 cell line as were those for viruses produced from transiently transfected HEK293T cells. Our procedure was thus shown to be highly efficient with regard to pseudovirus production for studies of the entry of SARS-CoV or SARS-CoV-2.

**Figure 4.**
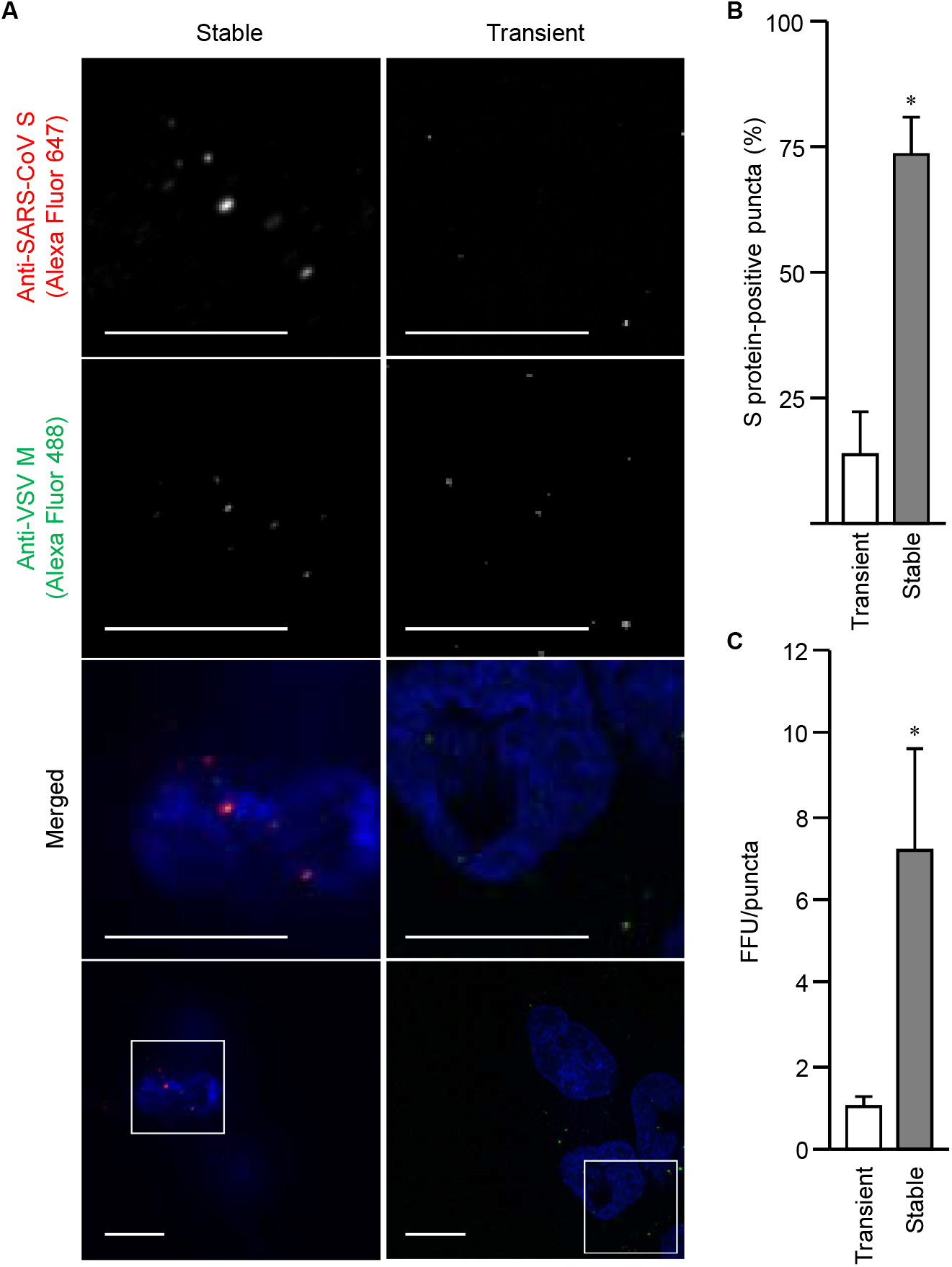
Production of pseudoviruses loaded with S protein of SARS-CoV-2 by a stable cell line. **(A)** BEAS-2B cells expressing ACE2 were exposed for 1 h at 4°C to pseudotyped viruses produced from a VeroE6 cell line stably expressing or HEK293T cells transiently expressing S protein of SARS-CoV-2. The cells were then fixed, stained with Hoechst 33342 (blue), subjected to immunofluorescence staining with antibodies to SARS-CoV S protein and VSV M protein, and examined with a confocal microscope. Representative images are shown. The boxed regions in the bottom panels are shown at higher magnification in the panels above. Bars, 10 μm. **(B)** Quantification of the fraction of S protein-positive puncta among all M protein–positive puncta in images as in (**A**). Data are means + SEM from three independent experiments. \**p* < 0.001 (Student’s t test). **(C)** FFU normalized by the area of puncta for the pseudoviruses produced as in (**A**) was determined as in **Figure S4**. Data are means + SEM from three independent experiments. *, *p* < 0.004 (one-way ANOVA with Tukey’s HSD post-hoc test). See also **Figure S4**.

In this study, we unexpectedly found that a large proportion of pseudotyped virus particles produced by the conventional protocol (Hoffmann et al., 2020; Rentsch and Zimmer, 2011) did not bear coronavirus S proteins. This finding raised the concern that S protein–free virions might affect the entry process of, or cellular responses to, S protein–bearing particles and thereby confound experimental outcomes. We therefore established a modified protocol based on stable cell lines in order to generate pseudoviruses for which a large proportion of the virus particles harbor heterologous envelope proteins. Alternative methods that are free of such issues relating to virus particles lacking envelope protein, including reverse genetic systems for in vitro virus assembly from SARS-CoV-2 full-length cDNA (Torii et al., 2021; Xie et al., 2020), have been developed. However, given that the VSV-based pseudotyped virus systems have been adopted by many researchers, the modification described here should be readily incorporated into current protocols and lead both to a better understanding of virus entry and to accelerated vaccine and therapeutic development for SARS-CoV-2.

## ACKNOWLEDGEMENTS

We thank Dr. S. Pöhlmann for expression plasmids, viruses and cells, as well as A. Kikuchi for technical assistance. This work was supported in part by Grants-in-Aid from the Ministry of Education, Culture, Sports, Science and Technology of Japan (#26115701 and #15H01248), from the Japan Society for the Promotion of Science (#19H05411, #20H05872, #21H02735 and #21H00413), and from the Japan Agency for Medical Research and Development (JP17fk0108124j0601), as well as by COVID-19 Drug and Vaccine Development Donation.

## AUTHOR CONTRIBUTIONS

Conceptualization, Y.F. and Y.O.; Methodology, Y.F. and S.K.; Formal analysis, Y.F. and S.K.; Investigation, Y.F., S.K., A.Y., A.O.S., and M.F.; Resources, Y.F., S.K., A.Y., A.O.S., and M.F.; Writing–original draft, Y.F. and Y.O.; Writing–review and editing, Y.F., M.A., Y.Y., and Y.O.; Visualization, Y.F. and Y.O.; Supervision, Y.Y. and Y.O.; Project administration, Y.O.; Funding acquisition, Y.F. and Y.O.

## STAR Methods

### Contact for reagent and resource sharing

Further information and requests for resources and reagents should be directed to and will be fulfilled by the Lead Contact, Yusuke Ohba.

### Cell lines

HEK293T (ATCC CRL-11268), VeroE6 (CRL-1586), and BEAS-2B (CRL-9609) cells were obtained from American Type Culture Collection (ATCC; Manassas, VA) and were maintained under a humidified atmosphere of 5% CO_2_ at 37°C in Dulbecco’s modified Eagle’s medium (DMEM) (Sigma-Aldrich, Tokyo, Japan) supplemented with 10% fetal bovine serum (FBS) (Capricorn Scientific, Ebsdorfergrund, Germany). BHK21/G43 were kindly provided by S. Pöhlmann (Leibniz Institute for Primate Research, Göttingen, Germany). Expression plasmids were introduced into cells by transfection for 24 h with the use of Polyethylenimine “Max” (PEI MAX; Polysciences, Warrington, PA). The absence of mycoplasma contamination was confirmed with the use of a PCR Mycoplasma Test Kit (Takara, Shiga, Japan).

### Reagents and antibodies

Antibodies to SARS-CoV S protein (#40150-R007), to RFP (#PM005), to VSV G protein (#EB0010), and to VSV M protein (#MABF2347) were obtained from Sino Biological (Beijing, China), Medical & Biological Laboratory (Nagoya, Japan), Kerafast (Boston, MA), and Merck (Darmstadt, Germany), respectively. Hoechst 33342, carbocyanine dyes (DiO, DiI, and DiD), and Alexa Fluor 647– or Alexa Fluor 488–labeled antibodies to rabbit or mouse immunoglobulin G were obtained from Thermo Fisher Scientific (Carlsbad, CA). Horseradish peroxidase (HRP)–conjugated goat secondary antibodies were obtained from Jackson ImmunoResearch Laboratories (West Grove, PA). Trypsin was obtained from FujiFilm-Wako (Osaka, Japan), and cell-dissociation solutions were from Thermo Fisher Scientific and Biological Industries (Beit-Haemek, Israel). Mifepristone was obtained from Sigma-Aldrich.

### Plasmids

Expression vectors for the S protein of SARS-CoV or SARS-CoV-2 (pCG1-SARS-CoV-S and pCG1-SARS-CoV-2-S) (Hoffmann et al., 2020) were kindly provided by S. Pöhlmann. The coding sequence for mCherry was cleaved out of pFX-Tom20-mCherry (Kashiwagi et al., 2019a) by digestion with *Bam*HI and *Bgl*II, and was then subcloned into the *Bam*HI sites of the expression vectors for the S proteins to obtain pCG1-mCherry-SARS-CoV-S and pCG1-mCherry-SARS-CoV-2-S. The pBABE-puro plasmid encoding a puromycin resistance gene was obtained from Addgene.

### Generation of stable cell lines

pCG1-mCherry-SARS-CoV-S or pCG1-mCherry-SARS-CoV-2-S was linearized with *Puv*I and introduced together with pBABE-puro into HEK293T or VeroE6 cells by transfection for 24 h with the use of PEI MAX. The cells were then cultured in DMEM supplemented with 10% FBS and puromycin (10 µg/ml), and puromycin-resistant colonies were collectively harvested. Expression of the mCherry-tagged S proteins was confirmed by immunofluorescence analysis.

BEAS-2B cells stably expressing human ACE2 were generated by lentivirus-mediated gene transfer. HEK293T cells were thus transfected with pLVX-ACE2-IRES-BLD (Daly et al., 2020), pCAG-HIVgp, and pCMV-VSVG-RSV-Rev (Miyoshi et al., 1998) for 48 h, after which the culture supernatant was collected. BEAS-2B cells were exposed to the recombinant lentivirus–containing supernatant for 1 h at 35°C, cultured for 2 days at 35°C in DMEM supplemented with 10% FBS, and further cultured for 14 days in DMEM supplemented with 10% FBS and blasticidin (20 µg/ml) (Funakoshi, Tokyo, Japan). The resulting blasticidin-resistant colonies were collectively harvested, and the constituent cells were maintained in DMEM supplemented with 10% FBS and blasticidin (15 µg/ml).

### Preparation of VSV pseudoviruses

Pseudotyped viruses were generated according to an established protocol (Rentsch and Zimmer, 2011) or its modified version developed in the present study. In brief, for preparation of VSVΔG-G, BHK21/G43 cells (which stably express VSV G protein) were incubated with mifepristone (10 nM) for 6 h and then infected with VSVΔG at a multiplicity of infection (MOI) of 100,000 FFU per cell for 16 h at 35°C. For generation of VSVΔG-S particles, cells expressing S protein of SARS-CoV or SARS-CoV-2 (either stably or transiently) were exposed to VSVΔG-G at an MOI of 3 FFU per cell for 16 h at 35°C in the presence of antibodies to VSV G in order to neutralize the parental viruses. The culture supernatants were passed through a filter (pore size of 0.45 µm) and subjected to ultracentrifugation, and the virus pellets were resuspended in phenol red-free DMEM/F12 (Thermo Fisher Scientific) and stored at –80°C until use. Given that the enhanced GFP (EGFP) gene is packaged in the genome of the pseudoviruses in this system (Rentsch and Zimmer, 2011), the titer of infectious viruses in culture supernatants can be determined by counting GFP-positive cells with use of a fluorescence microscope.

### Determination of FFU

BEAS-2B cells expressing ACE2 were incubated with pseudotyped viruses for 2 h at 35°C, and the culture supernatants were then removed and the cells cultured in DMEM for 16 h at 35°C before staining with Hoechst 33342 for 15 min and examination with a fluorescence microscope. The FFU value of the pseudovirus suspension was determined by counting the number of GFP-positive cells with the use of the “Multi-wavelength cell scoring” module of MetaMorph software (Molecular Devices, CA).

### Fluorescence microscopy

Fluorescence imaging and data analysis were performed essentially as described previously (Fujioka et al., 2019; Kashiwagi et al., 2019b). In brief, cells transfected with mCherry-CoV-S or mCherry-CoV-2-S vectors were stained with Hoechst 33342 for 15 min and placed in a stage-top incubation chamber maintained at 37°C on a Nikon Ti2 microscope (Nikon, Tokyo, Japan) equipped with a Zyla5.5 scientific complementary metal oxide semiconductor camera (Oxford Instruments, Belfast, UK). The cells were illuminated with an X-Cite turbo system (Excelitas Technologies, Waltham, MA) through a GFP HQ, Cy3 HQ, or DAPI-U HQ excitation filter (Nikon).

### Fluorescent labeling of pseudoviruses

For preparation of fluorescently labeled pseudoviruses, 1 ml of pseudovirus suspension was incubated for 1 h at room temperature with 6 µl of 100 μM DiD, DiI, or DiO stock solution in the dark (Fujioka et al., 2018; Nanbo et al., 2010). The stained particles were adsorbed to a PEI MAX–coated 96-well glass-based plate and observed with a confocal microscope.

### Immunofluorescence analysis

BEAS-2B cells expressing ACE2 were incubated with pseudotyped viruses for 1 h at 4°C to allow for virus attachment. The cells were then fixed with 3% paraformaldehyde for 15 min at room temperature and incubated with 1% bovine serum albumin to block nonspecific binding of antibodies. They were further incubated overnight at 4°C with primary antibodies (1:1000 dilution), after which immune complexes were detected by incubation for 1 h at room temperature in the dark with Alexa Fluor 488- or Alexa Fluor 647-conjugated secondary antibodies (1:250 dilution). Nuclei were visualized by staining with Hoechst 33342. Images were acquired with an IX83 microscope (Olympus, Tokyo, Japan) with X-lightV3 (CrestOptics, Rome, Italy).

### Immunoblot analysis

Transfected HEK293T cells were lysed in a lysis buffer [50 mM Tris-HCl (pH 7.4), 150 mM NaCl, 1%Nonidet P-40, 0.5% sodium deoxycholate, 0.1% SDS, 1 mM Na_3_VO_4_, cOmplete Protease Inhibitor Cocktail (Sigma-Aldrich)] for 30 min on ice. The lysates were centrifuged at 20,000 × *g* for 10 min at 4°C, and the resulting supernatants were subjected to SDS-polyacrylamide gel electrophoresis. The separated proteins were transferred to a polyvinylidene difluoride membrane (Bio-Rad, Hercules, CA) and subjected to immunoblot analysis. Immune complexes were detected with HRP-conjugated secondary antibodies, ECL Western Blotting Detection Reagent (Cytiva, Tokyo, Japan) and a MIIS imaging system (Givetechs, Sakura, Japan).

### Quantification and statistical analysis

Quantitative data are presented as means + SEM from at least three independent experiments and were compared with Student’s *t* test (parametric test between two conditions) or by one-way analysis of variance (ANOVA) followed by the Tukey honestly significant difference (HSD) post hoc test (among multiple conditions). No statistical methods were applied to predetermine sample size. Experiments were performed unblinded. A *p* value of <0.05 was considered statistically significant, and all statistical analysis was performed with JMP Pro software (version 15.0.0).

