## Supplementary material for "A method for the generation of pseudotyped virus particles bearing SARS coronavirus spike protein in high yields": sup

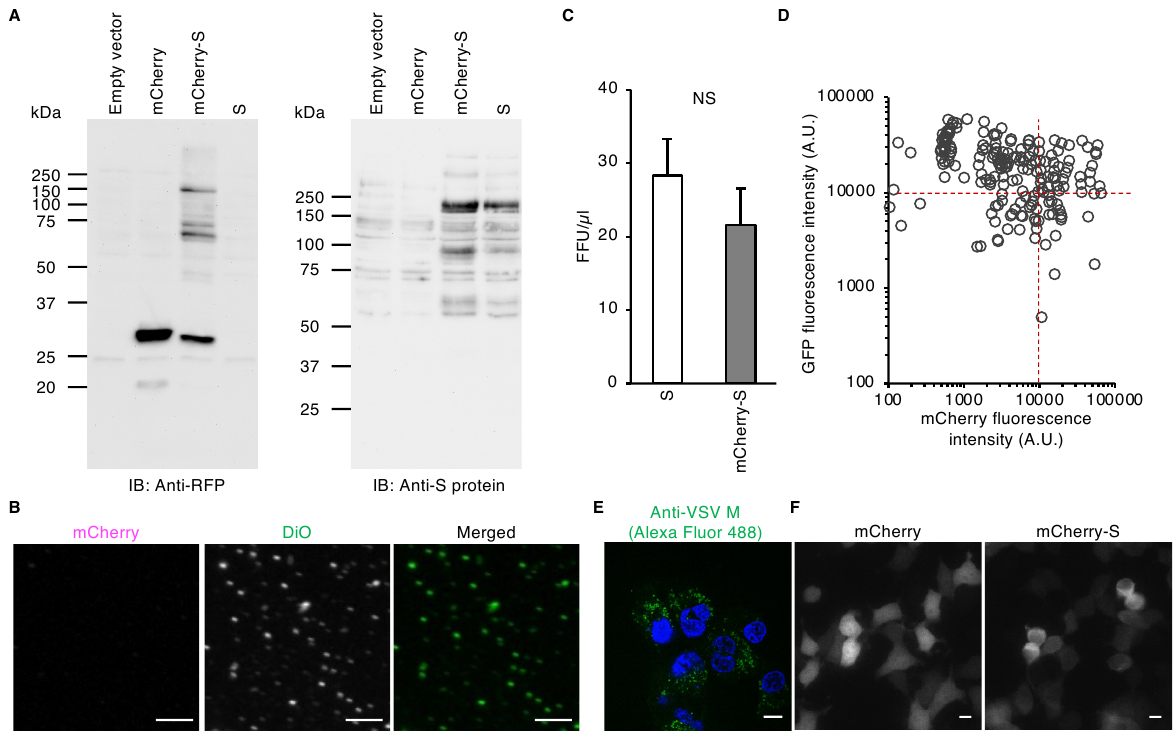

### Figure S1. Tagging of SARS-CoV S protein with mCherry does not affect pseudovirus production, related to Figure 1

(A) HEK293T cells transfected with an expression vector for mCherry, SARS-CoV S protein, or mCherry-tagged S protein (mCherry-S) for 24 h were lysed and subjected to immunoblot (IB) analysis with antibodies to RFP (for detection of mCherry) or to S protein. Representative blots are shown. Of note, mCherry is a derivative of an RFP and therefore detectable by an anti-RFP antibody.

(B) Pseudotyped viruses produced from HEK293T cells expressing mCherry-S and infected with VSVΔG-G were stained with DiO, allowed to adhere to a 96-well glass-based plate coated with polyethylenimine, and then observed with a confocal microscope for detection of mCherry and DiO fluorescence. Representative images are shown. Bars, 10 μm.

(C) BEAS-2B cells expressing ACE2 were exposed for 16 h to pseudotyped viruses produced from VSVΔG-G–infected HEK293T cells expressing S protein or mCherry-S protein. They were then stained with Hoechst 33342 and observed with a fluorescence microscope. The number of EGFP-positive cells (FFU), per microliter was determined. Data are means + SEM from three independent experiments. NS, not significant (Student’s *t* test).

(D) HEK293T cells transfected with an expression vector for mCherry and infected with VSVΔG-G for 16 h were observed with a fluorescence microscope, and the fluorescence intensities of EGFP and mCherry in individual cells were measured.

(E) BEAS-2B cells expressing ACE2 were exposed to pseudoviruses produced from empty vector-transfected, VSVΔG-G-infected HEK293T cells, fixed, and subjected to immunofluorescence analysis with antibodies to M protein of VSV (green fluorescence). Nuclei were stained with Hoechst 33342 (blue fluorescence). A representative image is shown. Bar, 10 μm.

(F) HEK293T cells expressing mCherry or mCherry-tagged S protein were observed with a fluorescence microscope. Representative images are shown. Bars, 10 μm.

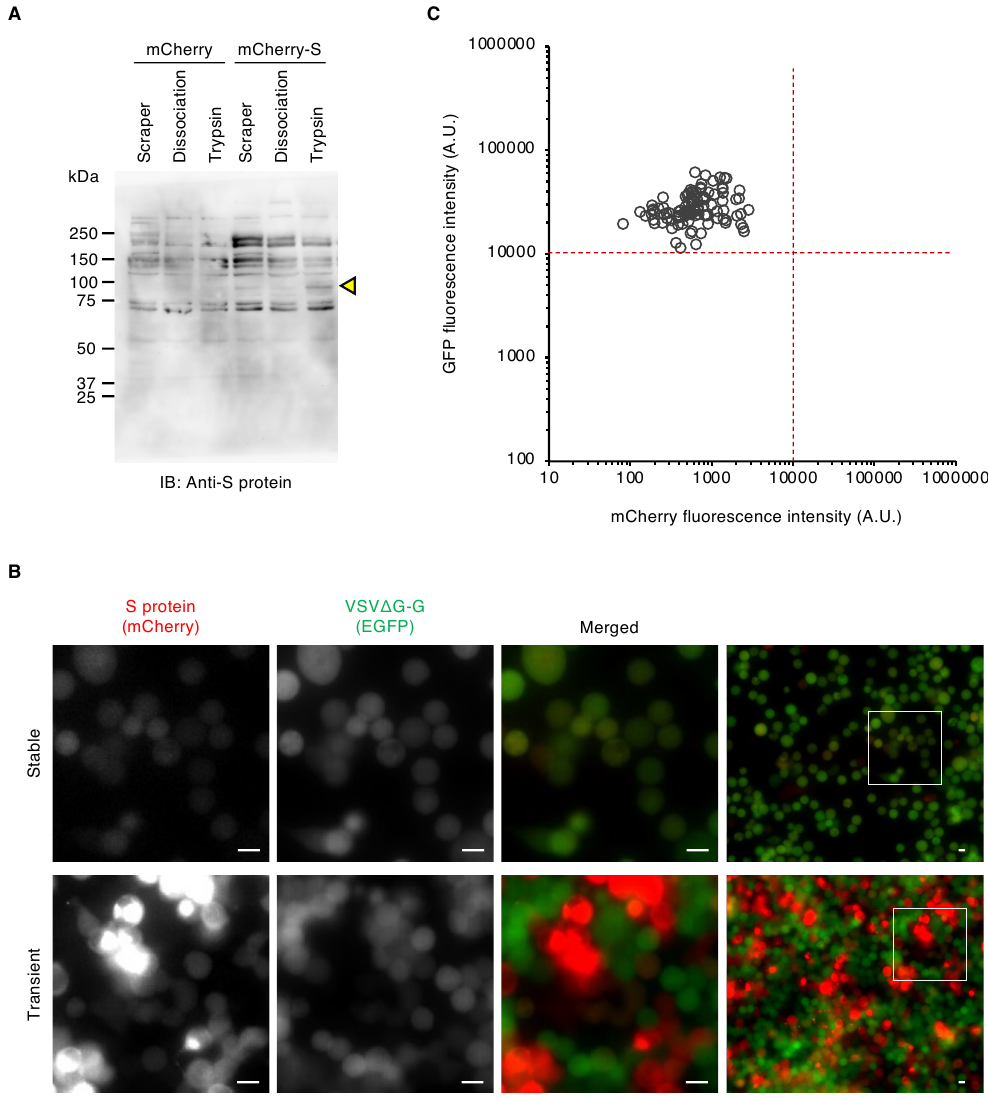

### Figure S2. VSVΔG-G can infect VeroE6 cells stably expressing SARS-CoV S protein, related to Figure 2

(A) VeroE6 cells expressing mCherry or mCherry-tagged SARS-CoV S protein were isolated either by exposure to trypsin or a cell-dissociation solution or with the use of a cell scraper. They were then lysed and subjected to immunoblot analysis with antibodies to SARS-CoV S protein. A representative blot is shown. The arrowhead indicates cleaved S protein.

(B) VeroE6 cells stably expressing or HEK293T cells transiently expressing mCherry-tagged S protein were exposed to VSVΔG-G for 16 h and then observed with a fluorescence microscope. Representative images of EGFP and mCherry fluorescence are shown. The boxed regions in the right panels are shown at higher magnification in the panels to the left. Bars, 10 μm.

(C) Scatter plot for the fluorescence intensities of EGFP and mCherry (mCherry-tagged S protein) in individual stably transfected VeroE6 cells treated as in (B).

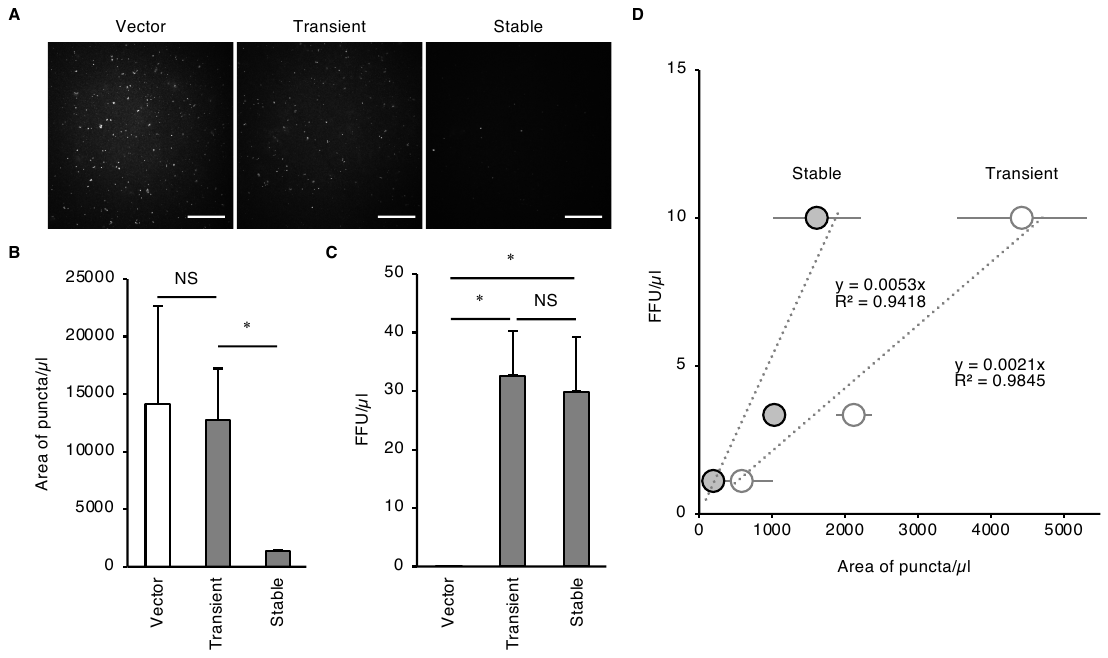

### Figure S3. Pseudotyped virus production from VeroE6 cells stably expressing SARS-CoV S protein, related to Figure 3

(A) Pseudotyped viruses produced either from HEK293T cells transiently transfected with an expression vector for SARS-CoV S protein (Transient) or with the corresponding empty vector (Vector) or from VeroE6 cells stably expressing SARS-CoV S protein (Stable) were stained with DiD, transferred to a polyethylenimine-coated glass-based plate, and observed with a confocal microscope. Representative images are shown. Bars, 10 μm.

(B) Quantification of the area of puncta for images as in (A). Data are means + SEM from three independent experiments NS, not significant; *, *p* < 0.001 (one-way ANOVA with Tukey’s HSD post-hoc test).

(C) ACE2-expressing BEAS-2B cells were exposed to pseudotyped viruses prepared as in (a) for determination of the number of GFP-positive cells and calculation of FFU per microliter of virus suspension. Data are means + SEM from three independent experiments NS, not significant; *, *p* < 0.001 (one-way ANOVA with Tukey’s HSD post-hoc test).

(D) DiD-labeled VSVΔG-S particles produced by the conventional (transiently transfected HEK293T cells) or modified (stably transfected VeroE6 cells) method were adsorbed onto polyethylenimine-coated plates at various dilutions and the area of puncta was measured with a confocal microscope and plotted against the corresponding FFU values determined as in (C). The regression equation and coefficient for each type of virus are indicated.

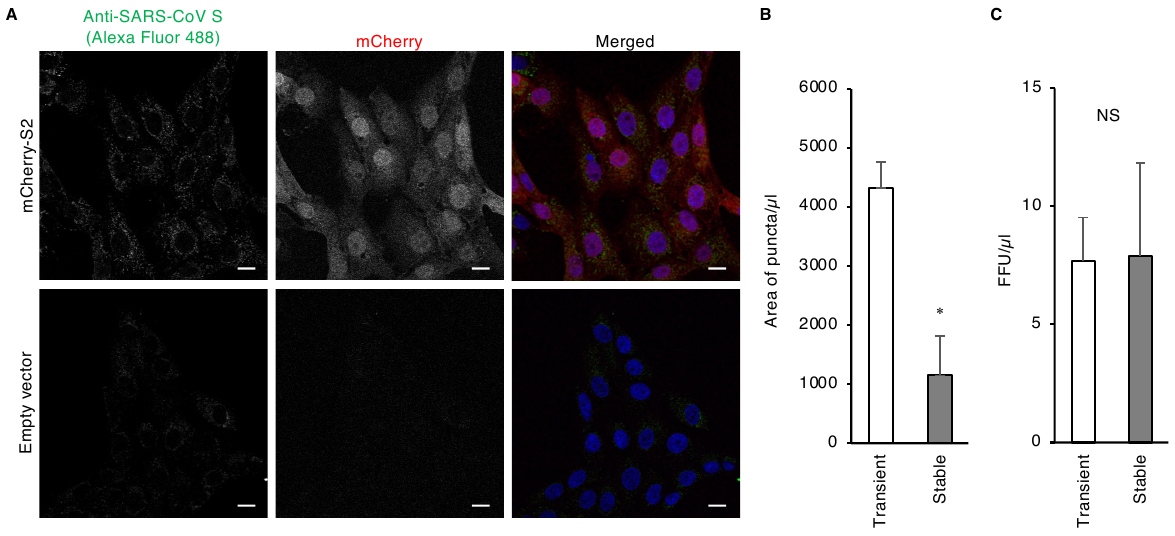

### Figure S4. Establishment of a VeroE6 cell line stably expressing S protein of SARS-CoV-2, related to Figure 4

(A) VeroE6 cells stably expressing mCherry-tagged S protein of SARS-CoV-2 (mCherry-S2) or parental VeroE6 cells were stained with Hoechst 33342, subjected to immunofluorescence staining with antibodies to SARS-CoV S protein, and examined by fluorescence microscopy. Representative images of S protein (green), Hoechst 33342 (blue), and mCherry (magenta) fluorescence are shown. Bars, 10 μm.

(B, C) The area of puncta (B) and FFU (C) per microliter of virus suspension were determined as in Figure S3 for VSVΔG-G pseudoviruses produced either from the cell line in (A) or from HEK293T cells transiently expressing SARS-CoV-2 S protein. Data are means + SEM from three independent experiments NS, not significant; *, *p* < 0.02 (Student’s *t* test).
